# Freshwater diatom biomonitoring through benthic kick-net metabarcoding

**DOI:** 10.1101/2020.05.25.115089

**Authors:** Victoria Carley Maitland, Chloe Victoria Robinson, Teresita M. Porter, Mehrdad Hajibabaei

## Abstract

Biomonitoring is an essential tool for assessing ecological conditions and informing management strategies. The application of DNA metabarcoding and high throughput sequencing has improved data quantity and resolution for biomonitoring of taxa such as macroinvertebrates, yet, there remains the need to optimise these methods for other taxonomic groups. Diatoms have a longstanding history in freshwater biomonitoring as bioindicators of water quality status. However, periphyton scraping, a common diatom sampling practice, is time-consuming and thus costly in terms of labour. This study examined whether the benthic kick-net technique used for macroinvertebrate biomonitoring could be applied to bulk-sample diatoms for metabarcoding. To test this approach, we collected samples using both conventional microhabitat periphyton scraping and bulk-tissue kick-net methodologies in parallel from replicated sites with different habitat status (good/fair). We found there was no significant difference in community assemblages between conventional periphyton scraping and kick-net methodologies, but there was significant difference between diatom communities depending on site quality (*P* = 0.029). These results show the diatom taxonomic coverage achieved through DNA metabarcoding of kick-net is suitable for ecological biomonitoring applications. The shift to a more robust sampling approach and capturing diatoms and macroinvertebrates in a single sampling event has the potential to significantly improve efficiency of biomonitoring programmes.

## Introduction

As climate change and other anthropogenic impacts continue to alter the environment, there is an increasing need for comprehensive ecological assessment. Rapid and robust biomonitoring is essential for informing management plans and mitigating further environmental degradation [1–3]. Freshwater biomonitoring typically involves sampling a range of aquatic taxa, with particular focus on biological indicator taxa, to assess environmental conditions based on diversity, richness, structure and function of the existing communities [3–5].

Traditionally, biomonitoring data is generated through morphological taxonomic classifications, however there has been a recent shift towards DNA-based identification using metabarcoding [6] coupled with high throughput sequencing [7]. In aquatic systems such as wadable streams, a combination of bulk-tissue benthic sampling using kick-net methodology with DNA metabarcoding, facilitates rapid data collection whilst maintaining data integrity [8–10]. The metabarcoding approach has been employed for numerous biomonitoring studies involving macroinvertebrates [11,12] for assessing freshwater health [5,10,13].

In addition to benthic macroinvertebrates, diatoms (members of Bacillariophyta) are also ideal biomonitoring target taxa for assessing freshwater system conditions [14–16]. These single-celled algae have a short generation time which allows for rapid responses to physical, chemical and biological changes in the environment [14,15,17]. Similar to macroinvertebrates, the high diversity and ubiquity of diatoms is used to create biotic indices that can accurately report freshwater quality [16,18,19]. Studies have shown that diatoms respond more readily to the presence of heavy metal pollutants compared to macroinvertebrates, which are generally more sensitive to shifts in hydrological conditions [17,20–22]. Monitoring only one of these taxonomic groups to assess overall ecosystem health could potentially cause gaps in knowledge that could subvert subsequent management strategies. Hence, diatoms are being used in a number of national and regional biomonitoring programmes.

Current methods for diatom sampling are time-consuming and laborious, which could hamper widespread use of diatoms for extensive freshwater biomonitoring [23,24]. The conventional diatom collection method involves the scraping of periphyton (a combination of algae, cyanobacteria, microbes, and detritus) from numerous substrates within littoral habitats [23–26]. These samples are then fixed and visualised using light microscopy [27–30]. From here, microscopy standards and keys are followed [29–31] to enable identification of diatoms to different taxonomic ranks. Within recent years, there has been the shift towards DNA metabarcoding-based identification of diatoms [15,16,32,33]. This involves the manual homogenized of periphyton scrapings into single samples, which are then processed via standard diatom metabarcoding procedures [34,35]. Alternative sampling methods, such as collection through the benthic kick-net technique, have not been tested for diatom biomonitoring applicability, however it is expected that this technique would drastically reduce time spent collecting samples. The ability to study diatom and macroinvertebrate assemblages from a single sample would allow biomonitoring programs to achieve an intensive appraisal of freshwater conditions. In a rapidly changing world, streamlining current methodology to obtain as much data in as little time as possible is crucial.

Because DNA-based analysis of environmental samples such as contents of a kick-net sample can provide a broad spectrum of organisms in the habitat sampled, we hypothesized that kick-net metabarcoding will provide diatom biodiversity comparable to commonly used scraping method. Specifically, we aimed to 1) investigate the feasibility of kick-net sampling for capturing community assemblages of freshwater diatoms versus conventional periphyton scraping using a high throughput sequencing coupled metabarcoding approach and 2) compare diatom community assemblages across a known habitat quality scale (Good and Fair) using both conventional and kick-net sampling to investigate presence of diatom indicator groups.

## Methods

### Field Sampling

Samples were collected in November 2019 from Grand River tributaries across four study sites in Waterloo, Ontario (Fig. 1). Status and location data were provided by Dougan & Associates based on a 2018 benthos biomonitoring project for the City of Waterloo (S1 Table). The four selected sites were a subset of the sites from this project and were chosen based on accessibility and habitat quality. Hilsenhoff Biotic Index ranges (weighted by species) informed the habitat quality scale [36] which categorized sites into ‘Good’ (4.51-5.50) and ‘Fair’ (5.51-6.50).

**Fig. 1.**
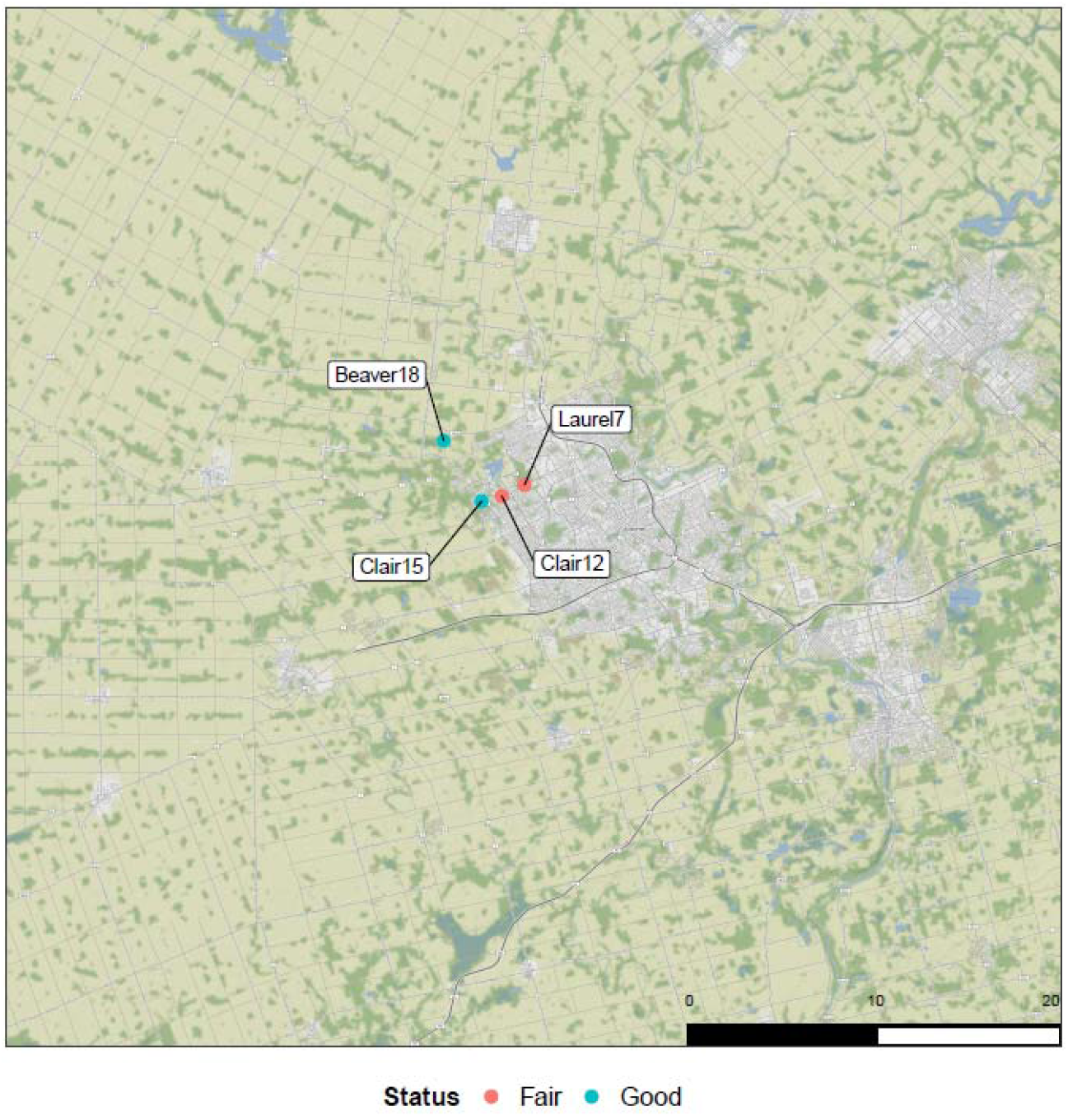
Map of sample sites located within the Waterloo region (Ontario, Canada) Scale bar shown in km, site habitat status indicated in legend.

Collection occurred in riffles, starting with a benthic kick-net sample, followed by subsequent periphyton scrapings of microhabitats representative of the reach (S2 Table). Periphyton scraping refers to the sampling of sediment, rock, macrophytes and leaf litter. Three replicates of each sampling type were collected at each site. Kick-net collection followed the Canadian Aquatic Biomonitoring Network [CABIN] protocol [37]. Effort was standardized to three minutes. The sampler moved up stream in a zig-zag pattern to encompass all microhabitats within the reach. Periphyton scraping samples were comprised of five specimens per microhabitat type to account for variability within the microhabitat [23]. Negative controls, consisting of molecular grade water, were collected prior to the collection of each rock sample (n= 9) to ensure the toothbrushes used for scraping biofilms from rocks had been adequately sterilised (S3 Table). All other samples were collected using manufacture-sealed sterile equipment. All samples were collected in 1L sample jars and placed in a cooler to transport back to the lab. Upon arrival at the lab, samples (n=45) were preserved using 100% ethanol and stored in a −20°C freezer until processing.

### Sample Validation and Extraction

To account for potential false negatives [38], diatom presence in the samples was confirmed using microscopy. A small amount of ethanol used to preserve the samples was placed on a slide and observed under a compound microscope at 100X magnification. Visual inspection confirmed the presence of diatoms in each sample type (S1 Fig.), however no taxonomic information was taken as morphological identification was beyond the scope of this study.

Once diatom presence was validated, samples were homogenized using standard blenders decontaminated by washing with ELIMINase® (VWR, Canada) then rinsing with deionized water before treating with UV light for 30 minutes. Homogenate was subsequently transferred to 50 mL Falcon tubes, where one tube was set aside and centrifuged at 2400 rpm for two minutes. Supernatant was removed and residual pellets were incubated at 70 °C until fully dried. Next, approximately 300 mg dried tissue was subsampled into PowerBead tubes and DNA extractions were completed using the DNeasy Power Soil kit (Qiagen, CA) following the manufacturer’s protocol. The only exception being that 50 μL of buffer C6 (TE) was used for final elution. Negative controls containing no tissue were also included with each batch of extractions. All negative controls failed to amplify and therefore were not sequenced.

### DNA Amplification, Library Preparation and Sequencing

Amplification targeted the 312 base pair long region of the chloroplast gene ribulose bisphosphate carboxylase large chain (rbcL) using five diatom specific primers. Following the methods of Rivera et al. [39], forward primers Diat_rbcL_708F_1 (5’- AGGTGAAG- TAAAAGGTTCWTACTTAAA-3’), Diat_rbcL_708F_2 (5’-AGGT- GAAGTTAAAGGTTCWTAYTTAAA-3’) and Diat_rbcL_708F_3 (5’-AGGTGAAAC- TAAAGGTTCWTACTTAAA-3’) were combined in an equimolar mix. Two reverse primers, Diat_rbcL_R3_1 (5’-CCTTCTAATTTACC- WACWACTG-3’) and Diat_rbcL_R3_2 (5’-CCTTCTAATTTACCWA-CAACAG-3’), were also combined and used for amplification. Each reaction used the following reagents: 17.5 μL HyPure™ molecular biology grade water, 2.5 μL 10X reaction buffer (200 mM Tris-HCl, 500 mM KCl, pH 8.4), 1 μL MgCl_2_ (50 mM), 05. μL dNTPs mix (10 mM), 0.5 μL of both forward (10 mM) and reserve (10 mM) equimolar mixes, 0.5 μL Invitrogen’s Platinum Taq polymerase (5 U) and 2 μL of DNA. Final reaction volume totaled 25 μL.

PCR protocol largely followed Rivera et al. [39] with minor adjustments. Instead of thirty cycles of denaturation at 95°C for 45 seconds, annealing at 55°C for 45 seconds and extension at 72°C for 45 seconds [39], this study increased the number of cycles to thirty-five. PCR amplification was also performed in two-steps, with the second PCR using 2 μL of amplicons from the first PCR instead of DNA, and Illumina-tailed primers. All PCRs were completed in Eppendorf Mastercycler ep gradient S thermal cycler. Successful amplification was confirmed using 1.5% agarose gel electrophoresis before purifying second PCR amplicons with the MinElute Purification kit (Qiagen). The next step was quantifying purified samples with a QuantIT PicoGreen daDNA assay kit and using these values to normalize all samples to 3 ng/μL. Samples were then indexed and pooled before purifying with AMpure magnetic beads. QuantIT PicoGreen daDNA assay kit was once again used to quantify the library and Bioanalyzer was used to determine fragment length. The library was diluted to 4 nM and 10% PhiX was added before being sequenced using Illumina MiSeq with a V3 MiSeq sequencing kit (300 × 2; MS-102-2003).

### Bioinformatic Processing

Illumina MiSeq paired-end reads were processed using the SCVURL rbcL metabarcode pipeline-1.0.2 pipeline available from https://github.com/terrimporter/SCVURL_rbcL_metabarcode_pipeline. SCVURL is an automated snakemake [40] bioinformatic pipeline that runs in a conda [41] environment. SeqPrep v1.3.2 [42] was used to pair raw reads requiring a minimum Phred score of 20 to ensure 99% base-calling accuracy. CUTADAPT v2.6 was used to trim primers from sequences, leaving a minimum fragment length of at least 150 base pairs [43]. Global exact sequence variant (ESV) [44] analysis was performed on the primer-trimmed reads. Reads were dereplicated using the ‘derep_fulllength’ command with the ‘sizein’ and ‘sizeout’ options of VSEARCH v2.14.1 [45]. VSEARCH was also used to denoise the data using the unoise3 algorithm [46]. These steps were taken to remove sequences with errors, chimeric sequences, PhiX carry-over and rare reads (singletons or doubletons) [47]. ESVs were classified using the rbcL diatom Classifier available from https://github.com/terrimporter/rbcLdiatomClassifier. Reference rbcL sequences were downloaded from the INRA diatom project [48]and reformatted to train the naive Bayesian classifier to make rapid, accurate taxonomic assignments [49]. This method makes assignments to the species rank and produces a statistical measure of confidence for each taxon up to the domain rank to help reduce false positive taxonomic assignments. We used 0.60 cutoff at the family rank (99% accuracy) and 0.20 cutoff at the genus rank (95% accuracy). The accuracy of the method assumes that target taxa are present in the reference database.

### Statistical Analysis

RStudio was used to analyze the data [50]. To account for variable reads within the library each sample was normalized to the 15^th^ percentile using the ‘rrarefy’ function in the vegan package [51,52].

ESV richness across the various sampling and status categories was calculated to assess differences between the methods and sites. A non-metric multi-dimensional (NMDS) analysis on Sorensen dissimilarities (binary Bray-Curtis) was conducted using the vegan ‘metaMDS’ function to determine if sampling method or site status created variation in community structure [5]. A scree plot was run using the ‘dimcheckMDS’ command from the goeveg package to determine the number of dimensions (k=2) to use with vegan metaMDS function[53]. Shephard’s curve and goodness of fit calculations were calculated using the vegan ‘stressplot’ and ‘goodness’ functions. The vegan ‘vegdist’ command was used to build a Sorensen dissimilarity matrix. We checked for heterogeneous distribution of dissimilarities using the ‘betadisper’ function. We used the ‘adonis’ function to perform a permutational analysis of variance (PERMANOVA). PERMANOVA was performed on conventional sampling methods (periphyton scraping) and kick-net methods, as well as site status to test for significant interactions between the categories [54].

To maintain a balanced design during statistical testing, we pooled all periphyton sampling into one sample type (conventional) and maintained kick-net samples as a separate sample type. The Jaccard index was calculated to assess the overall similarities between the sites, collection methods and site status. Nestedness and turnover of between kick-net and conventional samples were calculated using R package betapart function ‘beta-pair’ [55] followed by vegan function ‘betadisper’. The number of diatom family ESVs detected from kick-net or pooled conventional samples was also plotted. A dendrogram of diatom families detected was plotted using RAWGraphs (app.rawgraphs.io) and color-coded to show the samples the families were detected in [56]. Lastly, the frequency of ESVs detected from diatom families was visualized using a heatmap generated using geom_tile (ggplot) in R, plotting individual sample types for each site, split into two plots according to site status.

## Results

After bioinformatic processing, we generated 4,272 ESVs (2,166,157 reads). After taxonomic filtering (removal of non-diatom phyla), a total of 3,940 diatom ESVs (2,125,984 reads) were retained for data analysis. Read coverage per sample after normalisation (15^th^ percentile cut-off) was 37,735.

Since the rarefaction curves plateau, this indicated that the sequencing depth was sufficient to capture the ESV diversity in our PCRs (S2 Fig.). In terms of the top 10 orders identified, the order Naviculales represented 30.6% of ESVs (30% of reads) and Bacillariales represented 18.6% of ESVs (15.4% of reads; S3 Fig.).

### Taxonomic Coverage

In terms of taxonomic assignment, we identified a total of 1 phyla (Bacillariophyta), 4 classes, 23 orders, 44 families and 77 genera at the 95% correct assignment level. ESV richness varied across different sampling methods (Fig. 2). Mean overall ESV richness was used to calculate alpha diversity which displayed very similar values for all sampling methods across the four sites (S4 Table). Averaged across sites, kick-net samples produced the lowest mean ESV richness (225 ± 85), with sediment samples producing the highest ESV richness (317 ± 92).

**Fig. 2.**
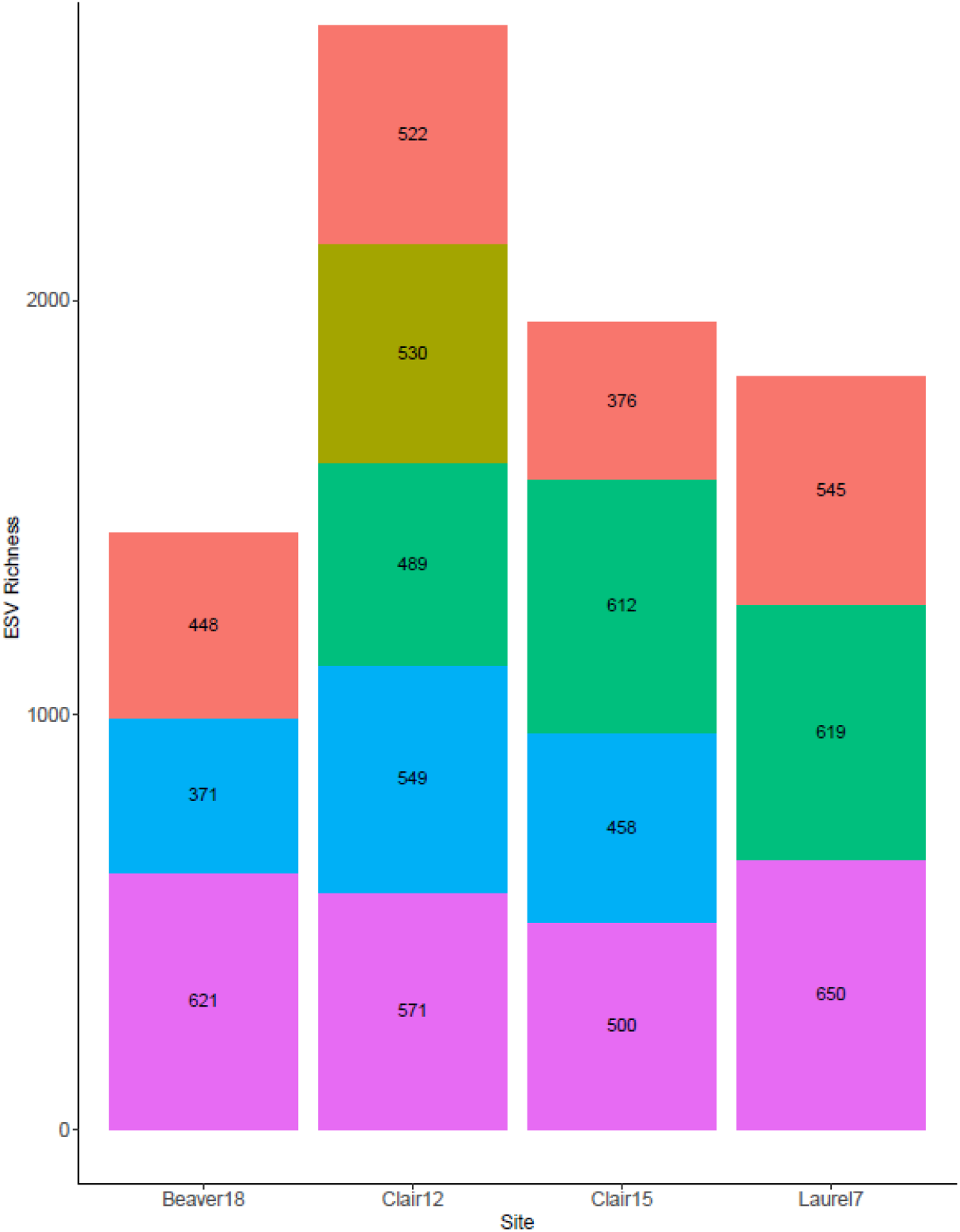
ESV richness varies across different sample types. Methods refer to the different sampling approaches analyzed (i.e. Kick-net, Macrophyte, Leaf Litter, Rock and Sediment). Replicates are pooled. Based on rarefied data.

Through investigating diatom families, a majority of families detected were present in all microhabitats and kick-net samples (Fig. 3). Two families (C oscinodiscaceae and Orthoseriaceae) were solely present in leaf litter samples and two families (Entomoneidaceae and Diadesmidaceae) were present only in sediment samples (Fig. 3).

**Fig. 3.**
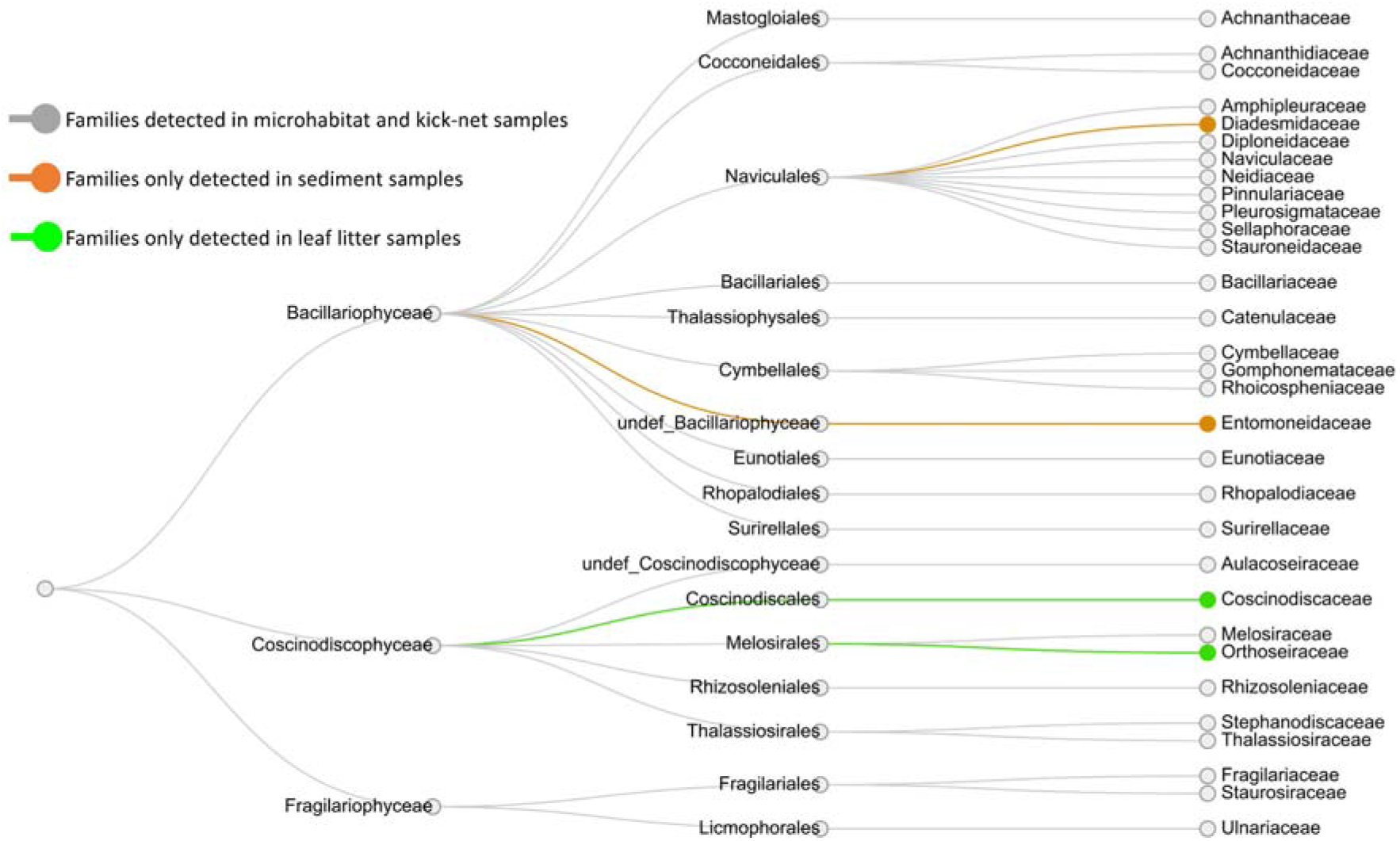
A majority of diatom families were detected in both microhabitat and kick-net samples.

In terms of diatom genera, some of the confidently identified genera represented by more than 2 sequence variants, identified from kick-net and conventional samples, included: *Nitzschia* (Bacillariales), *Polypedilum* (Chironomidae), *Navicula* (Naviculales), *Amphora* (Thalassiophysales) and *Ulnaria* (Licmophorales; Fig. 4).

**Fig. 4.**
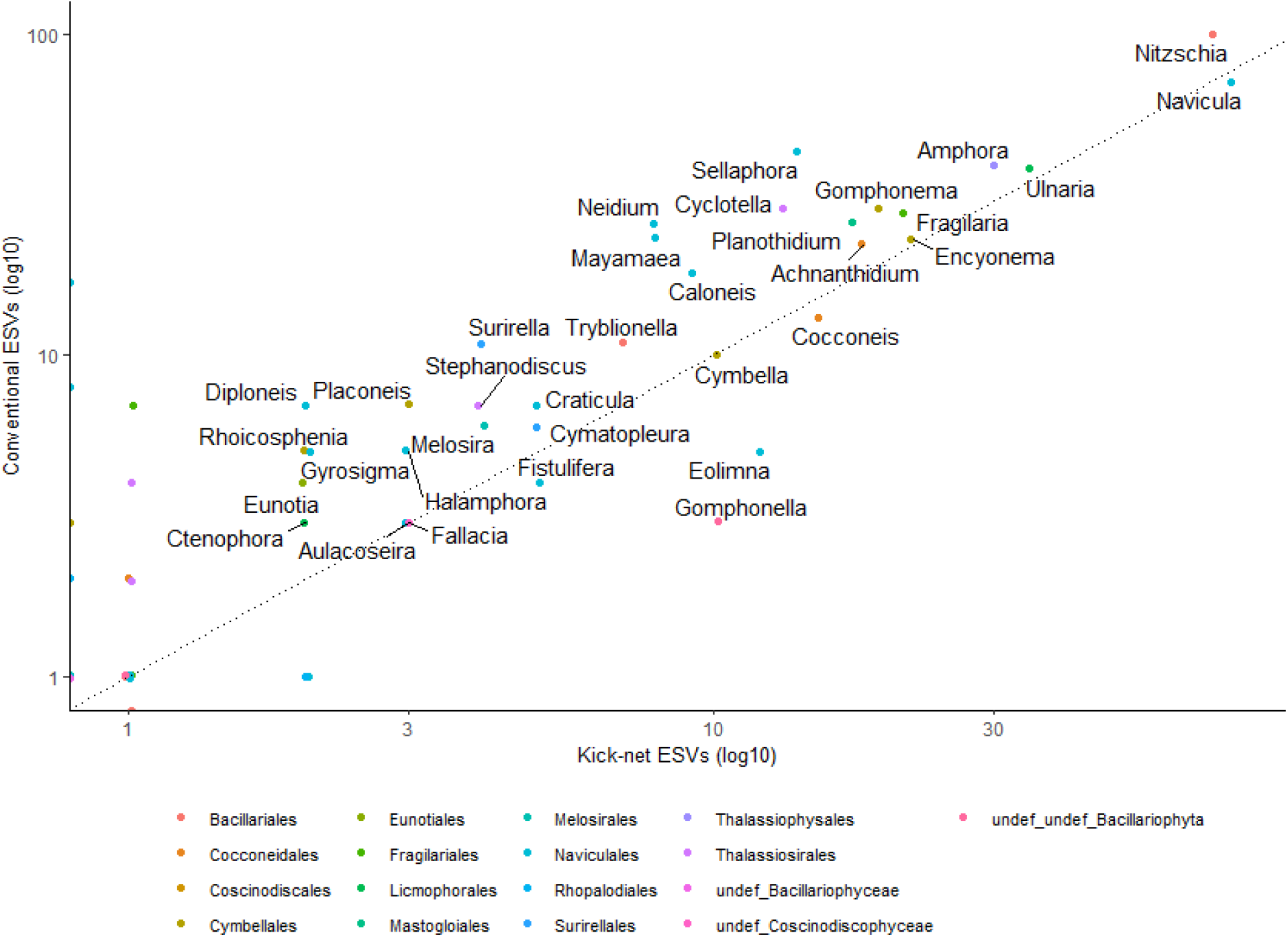
Number of ESVs detected from genera detected from kick-net versus conventionally sampled diatoms are similar. The points are color-coded for the orders detected in this study. A 1:1 correspondence line (dotted) is also shown. A log10 scale is shown on each axis to improve the spread of points with small values. Based on rarefied data.

### Diatom Diversity by Method and Site Status

NMDS plots showed that replicates clustered close together for site and status, with overlap observed between sampling methods and replicates (Fig. 5). When pooling conventional periphyton samples (i.e. macrophyte, leaf litter, rock, and sediment) at each site, there remained overlap between kick-net and conventional samples and samples also remained clustered by site and status (S4 Fig). PERMANOVA of the pooled samples, shows that analyzing data from kick-net or conventional samples (method) explains 13% of the variation in Bray Curtis dissimilarities (p-value = 0.776), sampling site (site) explains 58% of the variation (p-value = 0.009) and habitat quality status (status) explains 22% of the variation observed (p-value = 0.029; S5 Table). The Jaccard index for kick-net compared with conventional samples is 0.53, indicating samples are 53% similar, whereas the Jaccard index for fair compared to good site quality status samples is 0.20, indicating samples are only 20% similar. In terms of beta diversities of communities aggregated by the treatments of “kick-net” and “conventional”, there was no significant difference between turnover. For beta diversities of communities aggregated by site status, there was a significant difference between nestedness (*P* < 0.05) but not for turnover (*P* = 0.06). Fair samples appear to be significantly nested within good samples. These results further indicate that site status has a significant effect on the sampled community composition whereas conventional versus kick-net sampling methods do not.

**Fig. 5.**
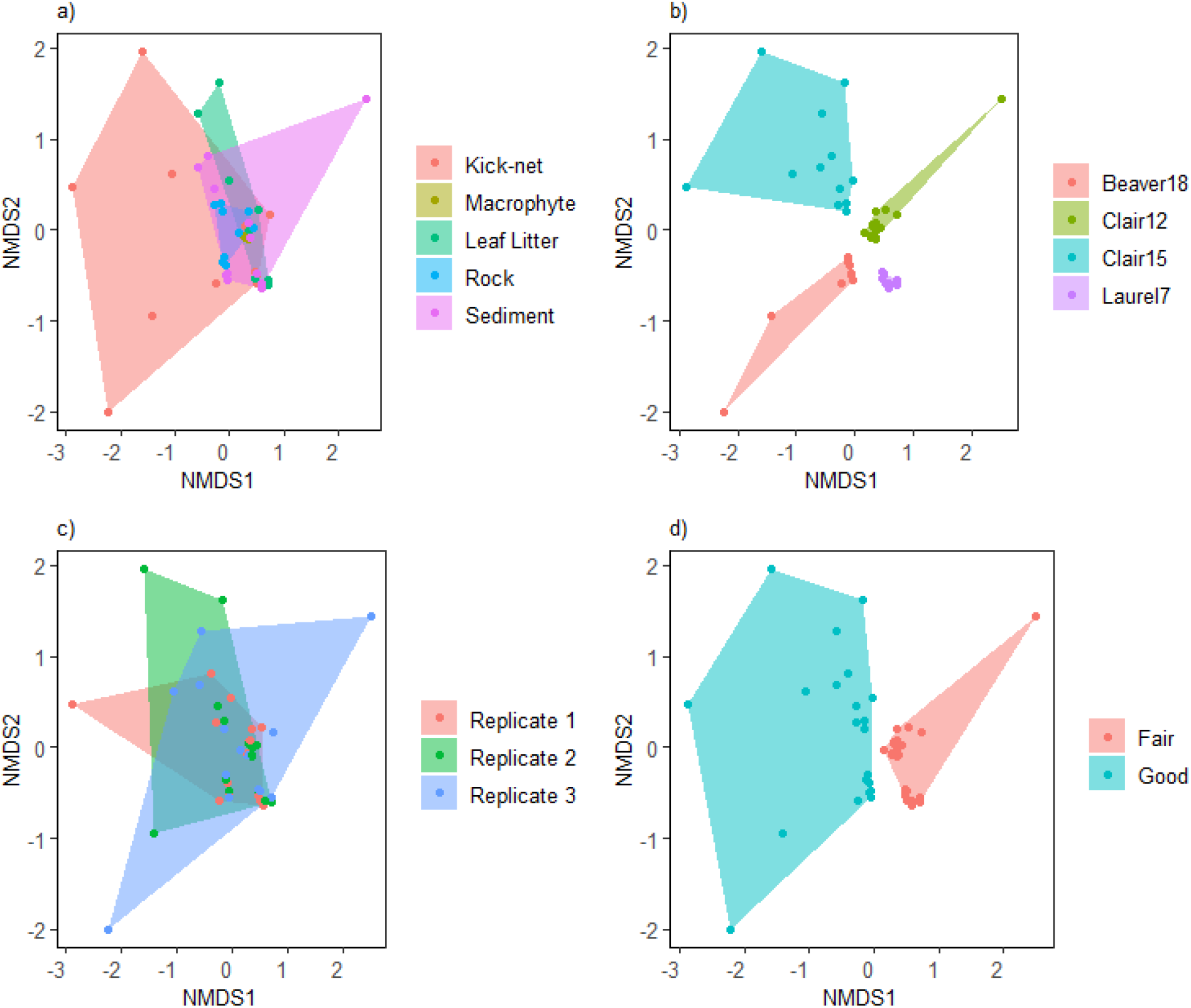
Non-metric multi-dimensional scaling plots show clustering mainly due to site and status. Specifically, a) binary Bray Curtis (Sorensen) dissimilarities overlapping across different sampling approaches, b) clustering by site, c) overlap between replicates, and d) clustering based on habitat quality status (stress = 0.111, R^2^ = 0.98). Based on rarefied data.

For individual sample types (i.e. kick-net, macrophyte, leaf litter, rock, and sediment), the heatmap shows that kick-net samples are largely representative of the diversity of families detected within each conventional periphyton sampling method (Fig.6.). In some cases, kick-net samples failed to detect diatom families which were present in conventional periphyton samples (e.g. Sellaphoraceae and Diadesmidaceae in Clair15) and conversely, kick-net samples also detected families which were not detected in conventional periphyton samples (e.g. Eunotiaceae and Neidiaceae in Clair12; Fig. 6). Similar assemblages of diatoms communities were detected across both fair and good quality sites, with the main difference observed between fair and good sites being the number of reads produced for families such as Thalassiosiraceae which was detected with a high number of reads (1000+) in fair sites and a lower number of reads (10-100) in good sites (Fig. 6).

**Fig. 6.**
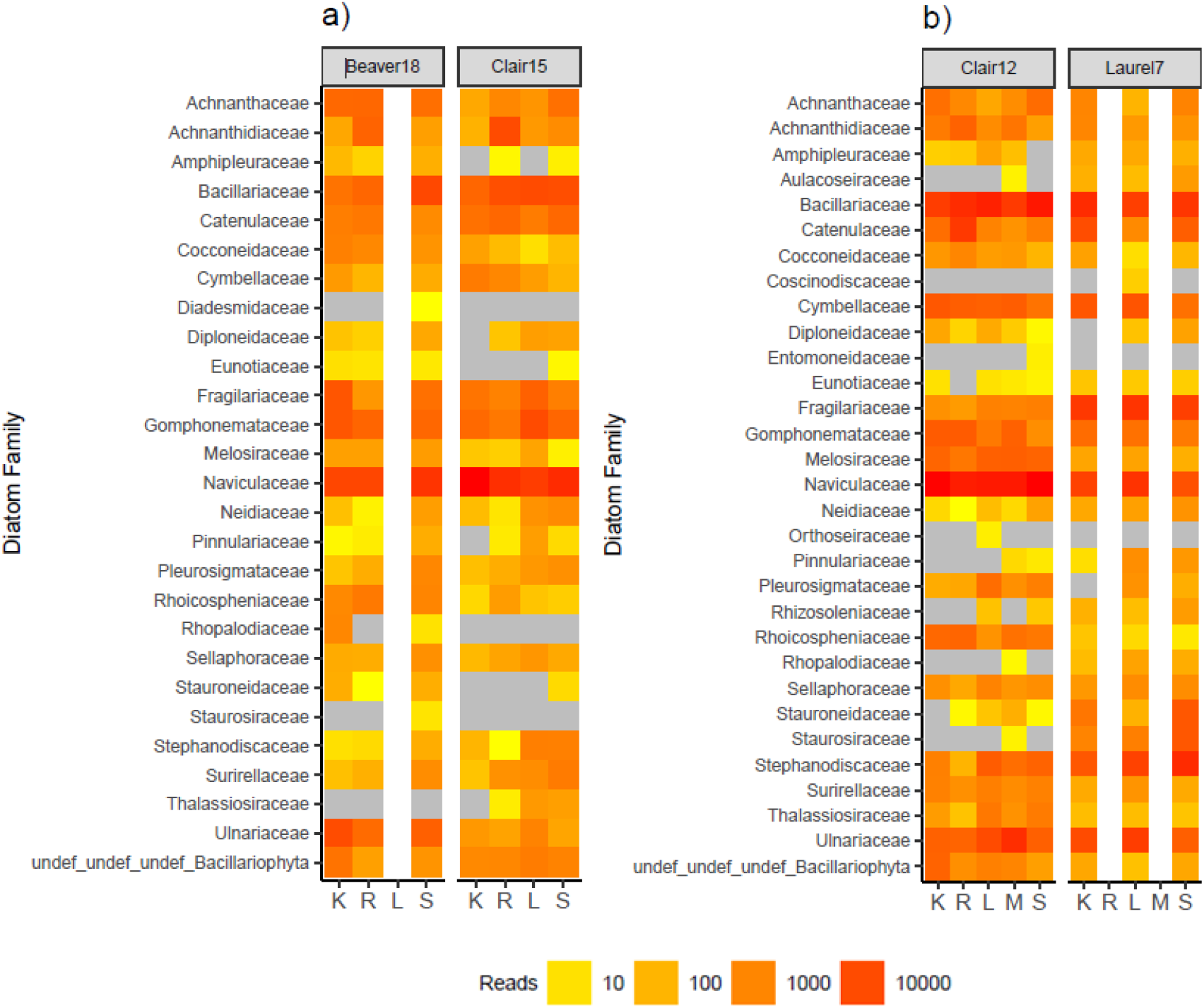
Samples detect similar diatom families across sampling methods and site status. Only ESVs taxonomically assigned to families with high confidence (bootstrap support >= 0.60 for 95% accuracy) are included. Part a) shows sites with a ‘good’ quality status b) sites with a ‘fair’ quality status. Sampling methods: K = kick-net; R = rock scraping; L = leaf litter; M = macrophyte; S = sediment. Empty lanes indicate the corresponding microhabitat was not present at the site. For each site, three replicates for each sampling method are pooled. Based on normalized data.

## Discussion

The demand for high-quality, reproducible ecological data is increasing in conjunction with the degradation of ecosystems globally [57]. There is a need to further streamline existing biomonitoring methodologies without sacrificing the quality of data produced [4,7,54,58]. With diatom assemblages providing a unique insight into the water quality status of lentic and lotic systems, fast-tracking diatom data collection for ecological assessments is a priority [39]. We have demonstrated that kick-net methodology with DNA metabarcoding provides sufficient taxonomic coverage to potentially be utilised as a for assessing diatom biodiversity in freshwater systems.

Kick-net sampling technique, whereby a zig-zag path is taken across the reach, provided sufficient representation of existing diatom community assemblages within site-specific microhabitats. Samples derived from the kick-net technique were highly comparable with conventional samples in terms of diatom taxa detected, despite the kick-net approach being more passive compared to direct periphyton scraping. Specific diatom taxa are known to have ecological preferences for different freshwater microhabitats [59,60]. For watershed-level health estimates, it is beneficial to be able to efficiently detect the diversity of diatom taxa present without directly sampling each microhabitat within a reach. We have demonstrated that kick-net methodology can sufficiently capture the existing diatom biodiversity, ground truthed by comparing assemblages detected with periphyton scrapings.

Ultimately, the detection of bioindicator species is a key variable to consider when comparing biomonitoring methods, as these taxa are pivotal for detecting subtle differences in freshwater health [3,5,14]. Naviculaceae contains diatom species sensitive to herbicide exposure, which is a family we observed in all sites and with all collection methods [61]. Additionally, the bioindicator family Stephanodiscaceae, (a known tolerant taxon) [62], has a higher read abundance in ‘Fair’ sites compared to ‘Good’ in both conventional and kick-net sample types. Despite the direct sampling approach of periphyton rock scraping, this methodology failed to detect this family at one of the sites where kick-net samples were successful at detecting this benthic family. Rock scrapings are commonly used as the sole collection method for diatoms [14,39,63,64], which suggests that the kick-net approach facilitates the detection of taxa which otherwise may be missed from conventional sampling.

## Conclusion

Overall, this study found that benthic kick-net methodology enables a robust and detailed assessment of freshwater diatom communities. This methodology is a scalable option for generating a holistic insight into the health of freshwater systems. The high similarity of diatom taxa detected between methods and significant differences between diatom communities detected in sites of differing habitat quality, demonstrates that this rapid method can provide accurate, fine-resolution taxonomic results. Future research should examine the duo-analyses approach of macroinvertebrate and diatom communities from a single kick-net sample, to determine reproducibility of multi-taxa targeting with this method. Additionally, future studies should consider exploring the use of multiple markers (i.e. rbcL cpDNA versus 18S rRNA gene), to address level of taxonomic resolution that can be obtained with these markers commonly used for diatom DNA barcoding.

## Supporting Information

**S1 Table. Information on study sites, including GPS coordinates and site status.**

**S2 Table. Outline of collections methods used in this study.** Samples for periphyton scraping were taken from a depth no greater than 1m (King et al. 2006).

**S3 Table. Summary table of decontamination and sterilisation procedures undertaken for the equipment in this study.**

**S4 Table. Mean ESV values (replicates pooled) for each sample type across the four sites.** Based on normalised data.

**S5 Table. rbcL exact sequence variants (ESVs) are not significantly different between sampling methods (kick-net versus conventional periphyton sampling).** No significant beta dispersion was detected within groups (method, site, status). Only significant difference detected was rbcL ESVs between sites and status. Summary of PERMANOVA results based on a Sorensen dissimilarity matrix of rbcL ESVs. Significant p-values are bolded.

**S1 Fig. Example of confirmation of diatom presence from preservative of kick-net sample.** Image: CBG Photography Group.

**S2 Fig. All samples show that ESV sampling reached saturation.** Samples were color-coded by site or method as shown in the legend. The vertical dashed line indicates the 15th percentile of sampling read depth, which is the number of reads that would be used in any future analysis based on normalized data.

**S3 Fig. Naviculales is the most abundant diatom order detected.** Results for the top 10 orders are shown with respect to proportion of ESVs and reads recovered. Based on raw unnormalized data.

**S4 Fig. Non-metric multi-dimensional scaling plots of microhabitat samples pooled show clustering by due to site and status.** Specifically, a) depicts overlap between the binary Bray Curtis (Sorensen) dissimilarities between different sampling approaches, b) sample site clustering c) clustering based on habitat quality status (stress = 0.012, R2 = 0.98). Based on rarefied data.

## Acknowledgements

We would like to extend thanks to Michael Wright for providing advice on laboratory procedures, Josip Rudar for assistance with bioinformatics and Genevieve Johnson for help collecting samples. We would also like to acknowledge Jessica Robinson, Jaclyn McKeown, Allison Brown and Monica Young from the Centre for Biodiversity Genomics’ Collections department for assisting with imaging and use of equipment. This study is funded by the Government of Canada through Genome Canada and Ontario Genomics. Teresita Porter was funded by the Government of Canada through the Genomics Research and Development Initiative (GRDI) Metagenomics-Based Ecosystem Biomonitoring Project (Ecobiomics).

## Author Contributions and Competing Interests

V.C.M. collected samples, conducted molecular and genomic analyses and wrote the manuscript with help from all authors, C.V.R and M.H. designed the study, C.V.R contributed to sampling, bioinformatic processing and statistical analyses, T.M.P trained the classifier, contributed to bioinformatic processing and advised on data analysis. All authors helped to write/edit the manuscript. The authors have declared that no competing interests exist.

## Data Availability

Raw sequences will be available from NCBI SRA on acceptance. The SCVURL rbcL metabarcode pipeline-1.0.2 is available from https://github.com/terrimporter/SCVURL_rbcL_metabarcode_pipeline and the rbcLdiatomClassifier v1 we used is available on GitHub at https://github.com/terrimporter/rbcLdiatomClassifier.

