## Supporting Information for "Freshwater diatom biomonitoring through benthic kick-net metabarcoding"

**S1 Table. Information on study sites, including GPS coordinates and site status.**

|  | Quality Status | Latitude | Longitude | Number of kick-net samples | Microhabitats Sampled |
| --- | --- | --- | --- | --- | --- |
| Beaver 18 | Good | 43.4920012 | -80.609783 | 3 | 2 (Rock and Sediment) |
| Clair 12 | Fair | 43.4654088 | -80.571321 | 3 | 4 (Rock, Leaf Litter, Macrophyte and Sediment) |
| Clair 15 | Good | 43.46290086 | -80.58467794 | 3 | 3 (Rock, Leaf Litter and Sediment) |
| Laurel 7 | Fair | 43.4707269 | -80.556274 | 3 | 2 (Leaf Litter and Sediment) |

**S2 Table. Outline of collections methods used in this study.** Samples for periphyton scraping were taken from a depth no greater than 1m [23].

| Sampling Type | Collection Method Description |
| --- | --- |
| Kick-net | 400 µm mesh net and frame attached to a pole was placed in water with the mouth facing upstream. Sampler kicked their feet to disturb benthos while moving upstream in a zig-zag pattern for a total of three minutes. If obstruction occurred, a timer was stopped until net was once again free, and kicking could recommence. When time elapsed, the net was lifted from the water and drained before placing contents into 1 L sample jar. |
| Sediment | The top 2 mm of sediment was suctioned using a 50 mL pipette until 10 mL of sample was collected. Contents were then dispensed into a 1 L sample jar and sampler then repeated these steps four more times before closing and storing sample jar. |
| Rocks | Sampler randomly selected 5 rocks from the reach, all approximately 10-20 cm in intermediate axis. Rocks were placed in a tub and transported to the stream bank where a minimum of 100 cm^2^ [27] of biofilm was scraped into the sampling jar using a previously sterilised toothbrush. 100% ethanol in a squirt bottle was also used to direct the scrapings into the 1 L sampling jar. |
| Macrophytes/Leaf Litter | Macrophytes and leaf litter samples were collected in an identical manner. Small handfuls of macrophytes representative of the reach and leaf litter was collected five times and placed into a 1 L sampling jar ¼ filled with water. Once all foliage was placed in the jar, lid was closed, and sample was shaken vigorously for 45 seconds. After time was complete, macrophytes and/or leaf litter was gently rubbed to remove any remaining periphyton before being discarded. Sample jar containing periphyton-water mixture was then closed and set aside. |

**S3 Table. Summary table of decontamination and sterilisation procedures undertaken for the equipment in this study.**

| Equipment | Decontamination/Sterilisation Procedure |
| --- | --- |
| Kick-nets | Nets were submerged in 10% bleach solution for 20 minutes then soaked in water for 20 minutes. When complete, nets were placed in clean garbage bags until use. |
| Sample jars | Sample jars were cleaned with ELIMINase® (VWR, Canada) and scrubbed with a soft bristle brush before being rinsed with deionized water. Once clean, jars were treated with UV light for 30 minutes then closed and sealed in plastic bags until use. |
| Toothbrushes | Toothbrushes were removed from original packaging and soaked in 10% bleach solution for 20 minutes, then water for an additional 20 minutes. Toothbrushes were then treated with UV light for 30 minutes and sealed in plastic bags until use. |

**S4 Table. Mean ESV richness for each sample type across the four sites.** Data was pooled across replicates. Based on normalised data.

|  | Beaver 18 | Clair 12 | Clair 15 | Laurel 7 | Average (± SD) |
| --- | --- | --- | --- | --- | --- |
| Kick-net | 174 | 262 | 138 | 326 | 225 (± 85) |
| Macrophyte | N/A | 331 | N/A | N/A | 331 (± 0) |
| Leaf litter | N/A | 275 | 255 | 356 | 295 (± 44) |
| Rock | 250 | 310 | 257 | N/A | 272 (± 27) |
| Sediment | 389 | 249 | 227 | 404 | 317 (± 92) |

N/A: sample not collected; SD: standard deviation.

**S5 Table. rbcL exact sequence variants (ESVs) are not significantly different between sampling methods (kick-net versus conventional periphyton sampling).** No significant beta dispersion was detected within groups (method, site, status). The only significant difference detected was rbcL ESVs between sites and status. Summary of PERMANOVA results based on a Sorensen dissimilarity matrix of rbcL ESVs. Significant p-values are bolded.

| Source of variation | Df | MS | F | R^2^ | P |
| --- | --- | --- | --- | --- | --- |
| A) sor ~ method | | | | | |
| Preservative | 1 | 0.264 | 0.890 | 0.130 | 0.776 |
| Residuals | 6 | 0.294 |  | 0.870 |  |
| Total | 7 |  |  | 1.000 |  |
| B) sor ~ site | | | | | |
| Site | 3 | 0.391 | 1.831 | 0.579 | **0.013*** |
| Residuals | 4 | 0.214 |  | 0.421 |  |
| Total | 7 |  |  | 1.000 |  |
| C) sor ~ status | | | | | |
| Status | 1 | 0.443 | 1.678 | 0.219 | **0.029*** |
| Residuals | 6 | 0.264 |  | 0.781 |  |
| Total | 7 |  |  | 1.000 |  |
| D) sor ~ method (strata = site) | | | | | |
| Method | 1 | 0.264 | 0.890 | 0.130 | 0.125 |
| Residuals | 6 | 0.294 |  | 0.870 |  |
| Total | 7 |  |  | 1.000 |  |
| E) sor ~ status (strata = site) | | | | | |
| Status | 1 | 0.443 | 1.678 | 0.219 | 0.056 |
| Residuals | 6 | 0.264 |  | 0.781 |  |
| Total | 7 |  |  | 1.000 |  |
| F) sor ~ status * method (strata = site) | | | | | |
| Status | 1 | 0.443 | 1.584 | 0.219 | 0.250 |
| Method | 1 | 0.264 | 0.945 | 0.130 | 0.125 |
| Status: Method | 1 | 0.201 | 0.718 | 0.090 | 0.500 |
| Residuals | 4 | 0.280 |  | 0.551 |  |
| Total | 7 |  |  | 1.000 |  |

sor (binary Bray Curtis sample by ESV matrix); Method (kick-net or conventional); Site (Clair 12, Clair 15, Beaver 18 or Laurel 07); Status (Fair or Good)

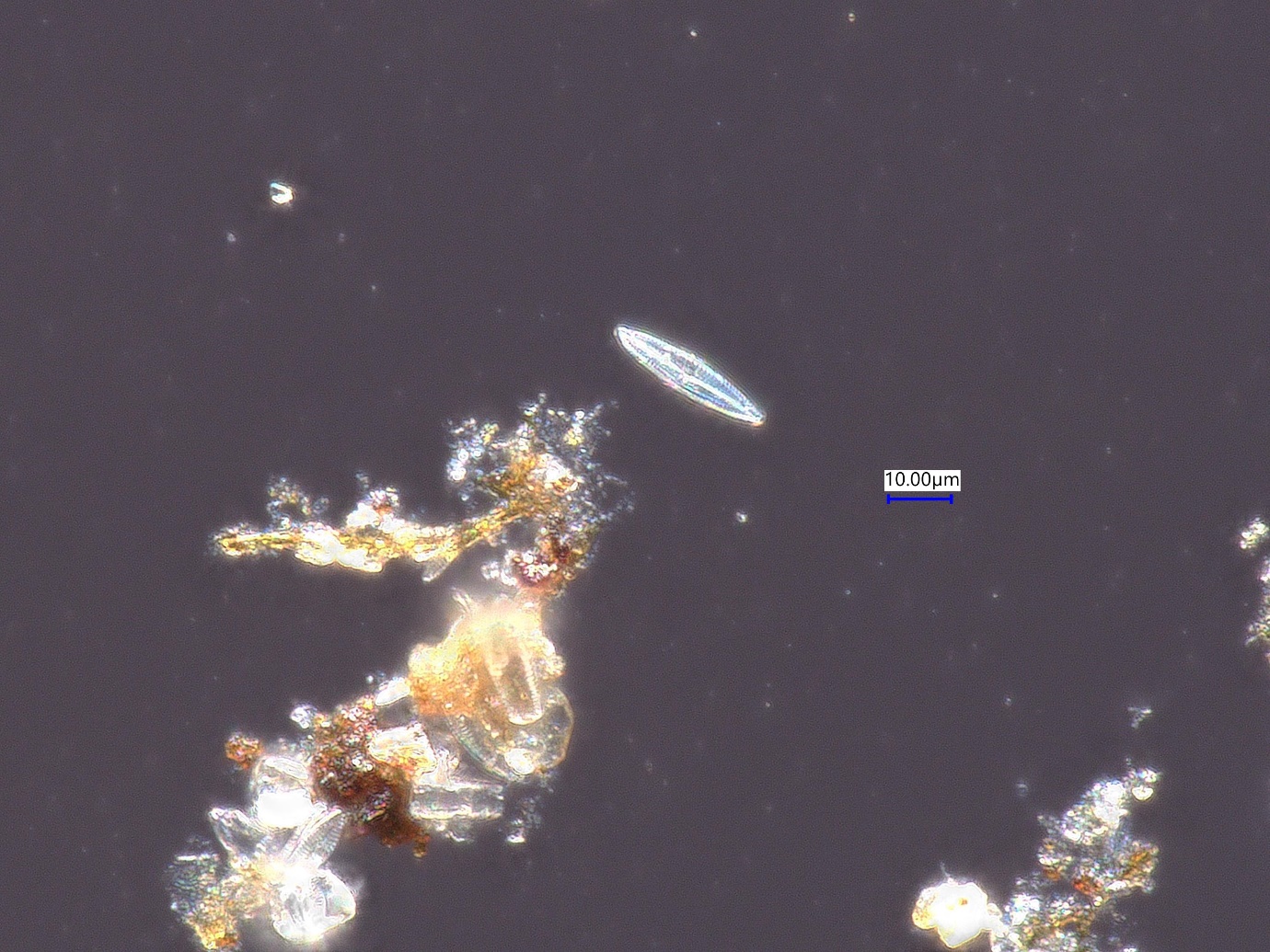

**S1 Fig. Example of confirmation of diatom presence from preservative of kick-net sample.** Image: CBG Photography Group

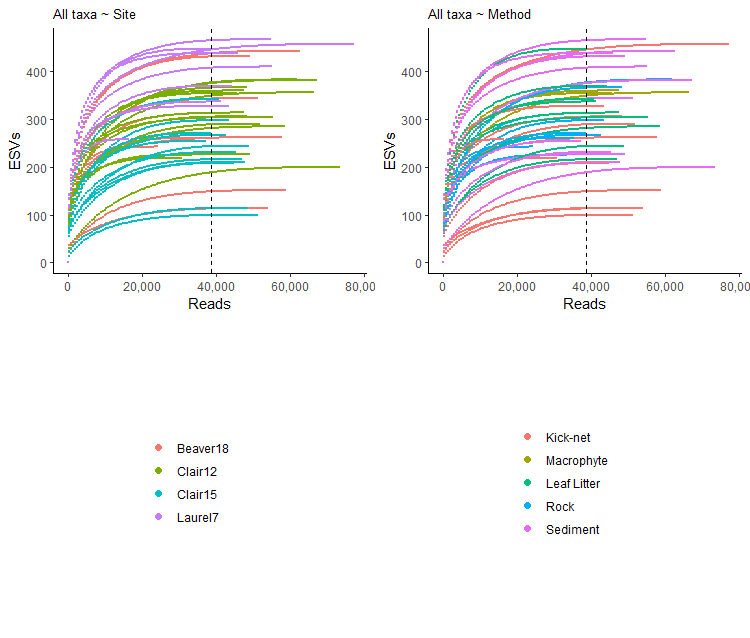

**S2 Fig. All samples show that ESV sampling reached saturation.** Each line represents reads from a sample plotted against the number of detected ESVs. Samples were color-coded by site or method as shown in the legend. The vertical dashed line indicates the 15th percentile of sampling read depth, which is the number of reads that would be used in any future analysis based on normalized data.

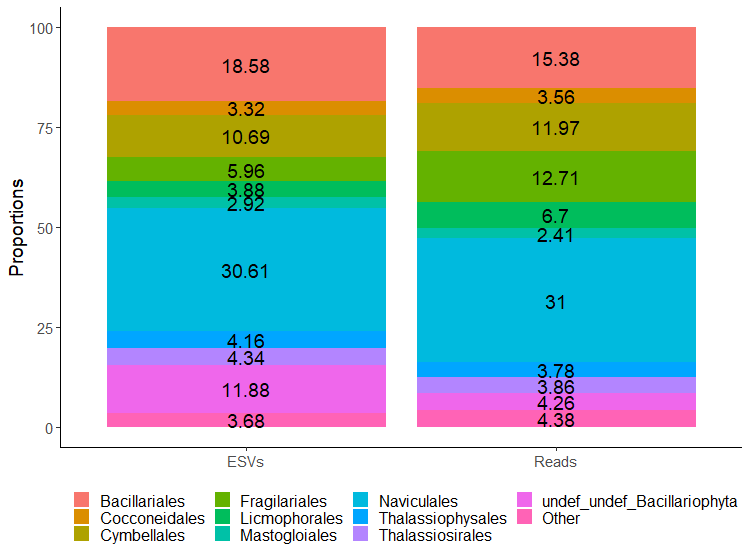

**S3 Fig. *Naviculales* is the most abundant diatom order detected.** Results for the top 10 orders are shown with respect to proportion of ESVs and reads recovered. Based on raw unnormalized data.

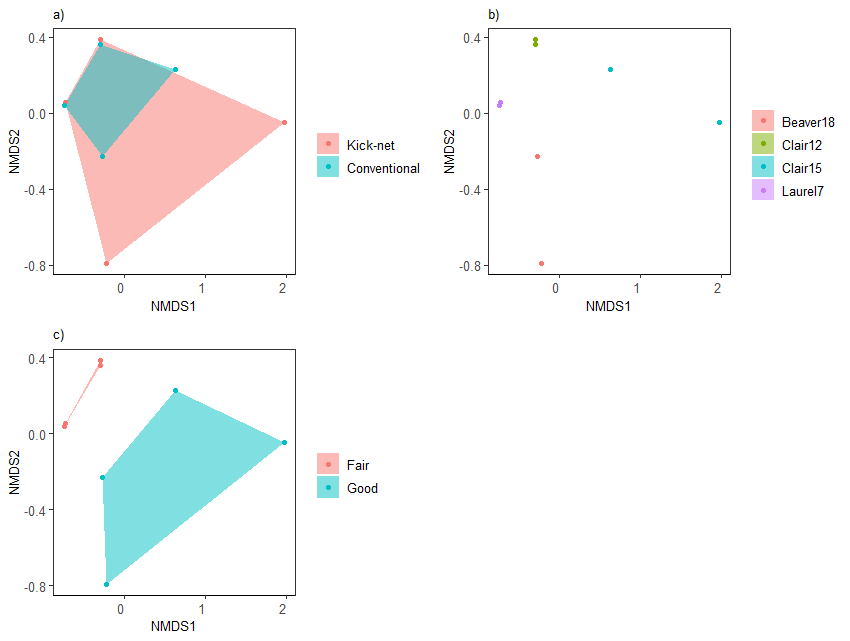

**S4 Fig.** **Non-metric multi-dimensional scaling plots of microhabitat samples pooled show clustering by due to site and status**. Specifically, a) depicts overlap between the binary Bray Curtis (Sorensen) dissimilarities between different sampling approaches, b) sample site clustering c) clustering based on habitat quality status. (stress = 0.012, R^2^ = 0.98). Based on rarefied data.
